# SERUM ANTIBODY RESPONSES AGAINST CARBAPENEM-RESISTANT *KLEBSIELLA PNEUMONIAE* IN INFECTED PATIENTS

**DOI:** 10.1101/2020.09.18.304469

**Authors:** Kasturi Banerjee, Michael P. Motley, Elizabeth Diago-Navarro, Bettina C. Fries

**Author notes:** **Corresponding Author: Bettina C. Fries**. **Authors Contributions:**, K.B., M.P.M, E.D.M. and B.C.F contributed equally to the ideation and development of the project. K.B. conducted all of the experiments, analyzed the data and generated all of the manuscript figures. K.B., M.P.M and B.C.F. contributed equally in writing and editing the manuscript. E.D.M. contributed in patient plasma and clinical isolates collection, and *wzi* typing. M.P.M. helped K.B in conducting animal experiments. **FinancialSupport:** This study was funded using US Veterans Affairs Merit Review Award 5I01 BX003741. BCF is an attending at the U.S. Department of Veterans Affairs - Northport VA Medical Center, Northport, NY. MPM is funded through F30 AI140611. The contents of this study do not represent the views of VA or the United States Government. **Meetings were the data was partially presented:** IDWeek 2019-IDSA, October 2 - 6, 2019, Washington, D.C., Abstract #: 405.

## Abstract

**Background:** Capsular polysaccharide (CPS) heterogeneity within carbapenem-resistant *Klebsiella pneumoniae* (CR-*Kp*) strain ST258 must be considered when developing CPS-based vaccines. We sought to characterize CPS-specific antibody responses elicited by CR-*Kp* infected patients.

**Method:** Plasma and bacterial isolates were collected from 33 hospital patients with positive CR-*Kp* culture. Isolates capsule were typed by *wzi* sequencing. Reactivity and measures of efficacy of patient antibodies were studied against 3 prevalent CR-*Kp* CPS types (*wzi29, 154* and *50*).

**Results:** High IgG titers against *wzi154* and *wzi50* CPS were documented in 79% of infected patients. Patient-derived (PD) IgGs agglutinated CR-*Kp* and limited growth better than naïve IgG, and promoted phagocytosis of strains across the serotype isolated from their donors. Additionally, poly-IgG from *wzi50 and wzi154* patients promoted phagocytosis of non-concordant CR-Kp serotypes. Such effects were lost when poly-IgG was depleted of CPS-specific IgG. Additionally, mice infected with *wzi50, wzi154, and wzi29* CR-*Kp* strains pre-opsonized with *wzi50* patient-derived IgG exhibited lower lung CFU than controls. Depletion of *wzi50* Abs reversed this effect in *wzi50 and wzi154* infections, whereas wzi154 Ab depletion reduced Poly-IgG efficacy against *wzi29* CR-*Kp*.

**Conclusions:** We are first to report cross-reactive properties of CPS-specific Abs from CR-*Kp* patients through both *in-vitro* and *in-vivo* models.

**Importance:** Carbapenem-resistant *Klebsiella pneumoniae* is a rapidly emerging public health threat that can cause fatal infections in ~40-50% of immunocompromised patients. Due to its resistance to nearly all antimicrobials, development of alternate therapies like antibodies and vaccines are urgently needed. Capsular polysaccharides are emerging as important immunotargets as they are a crucial for *Klebsiella pneumoniae* pathogenesis. Studying how host’s immune system protect against specific bacteria such as *Klebsiella pneumoiniae*, with respect to their specific capsule-type is crucial to develop effective anti-CPS immunotherapies. In this manuscript, we are the first to characterize humoral responses developed in infected patients against carbapenem-resistant *Klebsiella pneumoniae* expressing different *wzi*-capsule types. This study is the first to report efficacy of cross-reactive properties of CPS-specific Abs in both *in-vitro* and *in-vivo* models.

## Introduction

Infections with Carbapenem-resistant *Klebsiella pneumoniae* (CR-*Kp*) are associated with high mortality rates (2). Thus, the World Health Organization (WHO) considers the development of novel therapeutics for CR-*Kp* a critical priority (3). A large prospective observational multicenter study demonstrated that spread of CR-*Kp* In the United States is largely driven by expansion of sequence type (ST)258 or related clonal lineages (CG258) of CR-*Kp*. In that study CG258 represented 74% of carbapenemase-producing *Kp*(4). The genome of ST258 strains is highly conserved, with two dominant subclades (clade1 and clade2) varying only by several hundred kilobases, and *wzi* gene-based capsular typing has shown the predominance of only a few capsular polysaccharide subtypes (CPS) (5–7). Specifically, clade2 strains almost exclusively produce *wzi154* CPS, whereas whereas clade1 strains chiefly produce *wzi29* and *wzi50* capsules, but can produce other capsular types as well (8, 9). Importantly, infection with *wzi154/clade* 2 strains has been associated with worse prognoses than infections by other strains (9–12).

The polysaccharide capsule of *Kp* protects the pathogen from host cell-mediated killing and the bactericidal effect of serum, thus contributing to *Kp* pathogenicity (13). Studies examining the efficacy of antibodies (Abs) directed against CR-*Kp* CPS have demonstrated their potency in human cells and animal models (1, 14–17), and CPS-vaccines have the potential to prevent or moderate the severity of CR-*Kp* infection in monkeys, decrease the prevalence and emergence of resistant strains (14, 18–20). However, it is unknown if natural antibody responses to *K. pneumoniae* rely on anti-capsular responses. Given the heterogeneity of the CPS, it is important to understand whether diverse *wzi* CPS types are recognized by the immune system differentially during an infection. Furthermore, it is still unknown whether anti-CPS antibodies (Abs) cross-react and cross-protect across clades that express CPS with different *wzi*-types.

To understand whether vaccination with CPS constitutes a feasible strategy to elicit a protective immune response and mitigate CR-*Kp* emergence, we characterized the humoral response in a cohort of hospitalized patients infected or colonized with CR-*Kp*. We compared the Ab response elicited by different *wzi* capsule types (*wzi29, 154*, and *50*) and assessed the protective opsonophagocytic efficacy of human anti-CPS polyclonal IgG against CR-*Kp in vitro* and *in vivo*. This study is the first to our knowledge to document anti-CPS humoral responses in patients infected with different CR-*Kp* strains that produce specific *wzi*-type CPS. Our data indicates a cross-reactive therapeutic potential of the *wzi50* anti-CPS Abs for the first time. The implications of these findings for efforts of developing anti-capsular therapeutics are discussed in this study.

## Results

### Demographics and clinical variables

We identified 36 patients who were hospitalized at Stony Brook University Hospital between 2017-2019 with bacterial cultures that grew CR-*Kp*. Plasma samples were obtained from 33, while 3 did not give consent. Of those, 23 met published criteria for “symptomatic infection with CR-*Kp*”, while 10 were classified as “asymptomatic infection or colonized with CR-*Kp*” (12). Both cohorts (infected and colonized with CR-*Kp*) were similar in age and gender distribution, but infected patients had median length of stay (LOS) of 20.5 days (IQR= 14-52.5 days P<0.0001), whereas colonized patients had median LOS of 17 days (IQR= 8-33 days, P=0.0003). Subsequently, the median time to first positive culture from the day of admission was longer for infected patients compared to colonized patients (7.5 vs 3 days, respectively). Capsular serotyping by *wzi* sequencing showed 12 isolates to be *wzi154*, 10 to be *wzi29*, 2 to be *wzi50*, and 9 to be of other *wzi* types **(Table 1)**. *wzi154, 29*, and *50* were previously identified to be the most common *wzi* types among ST258 isolates in the New York City Area (8).

**Table I:**
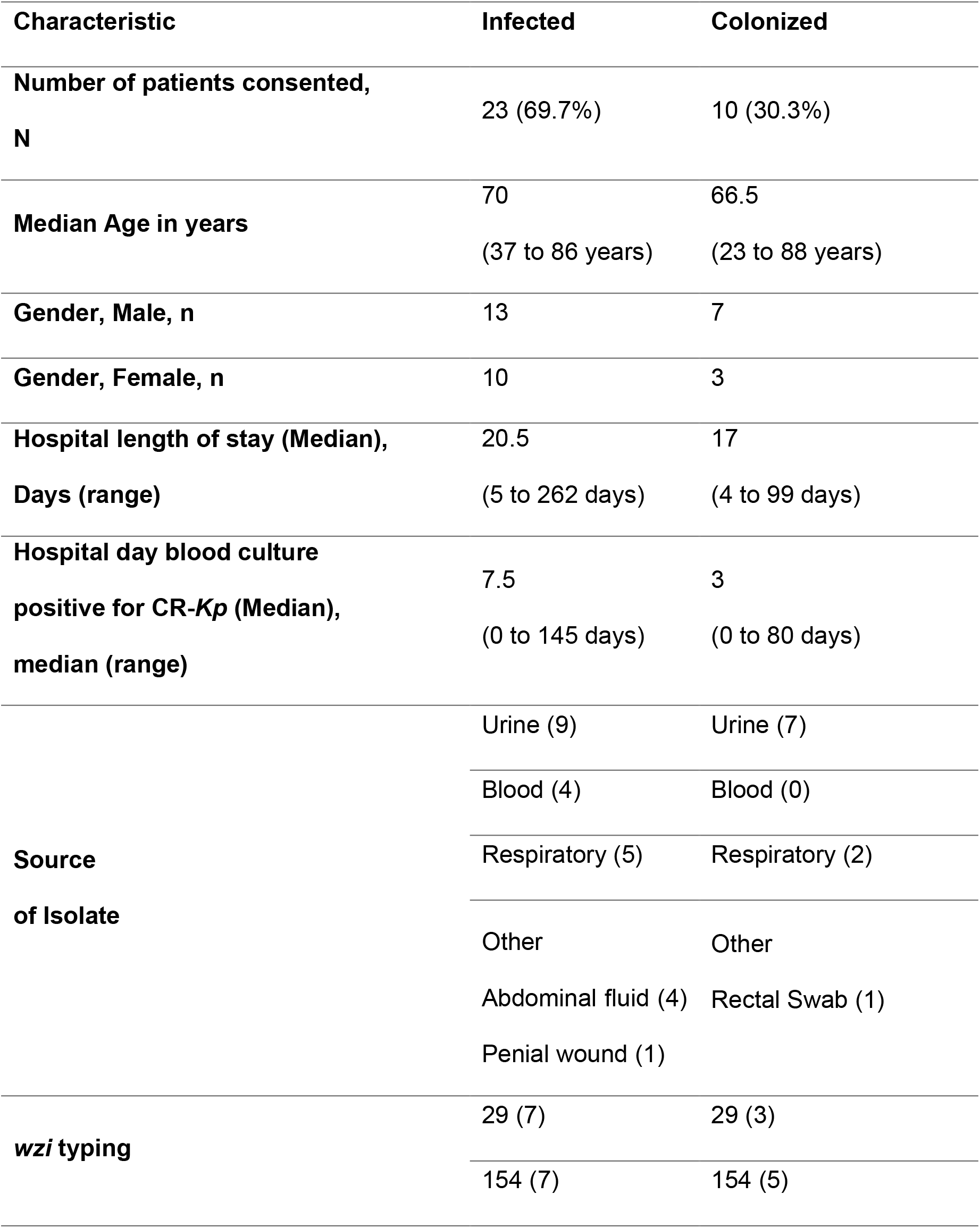
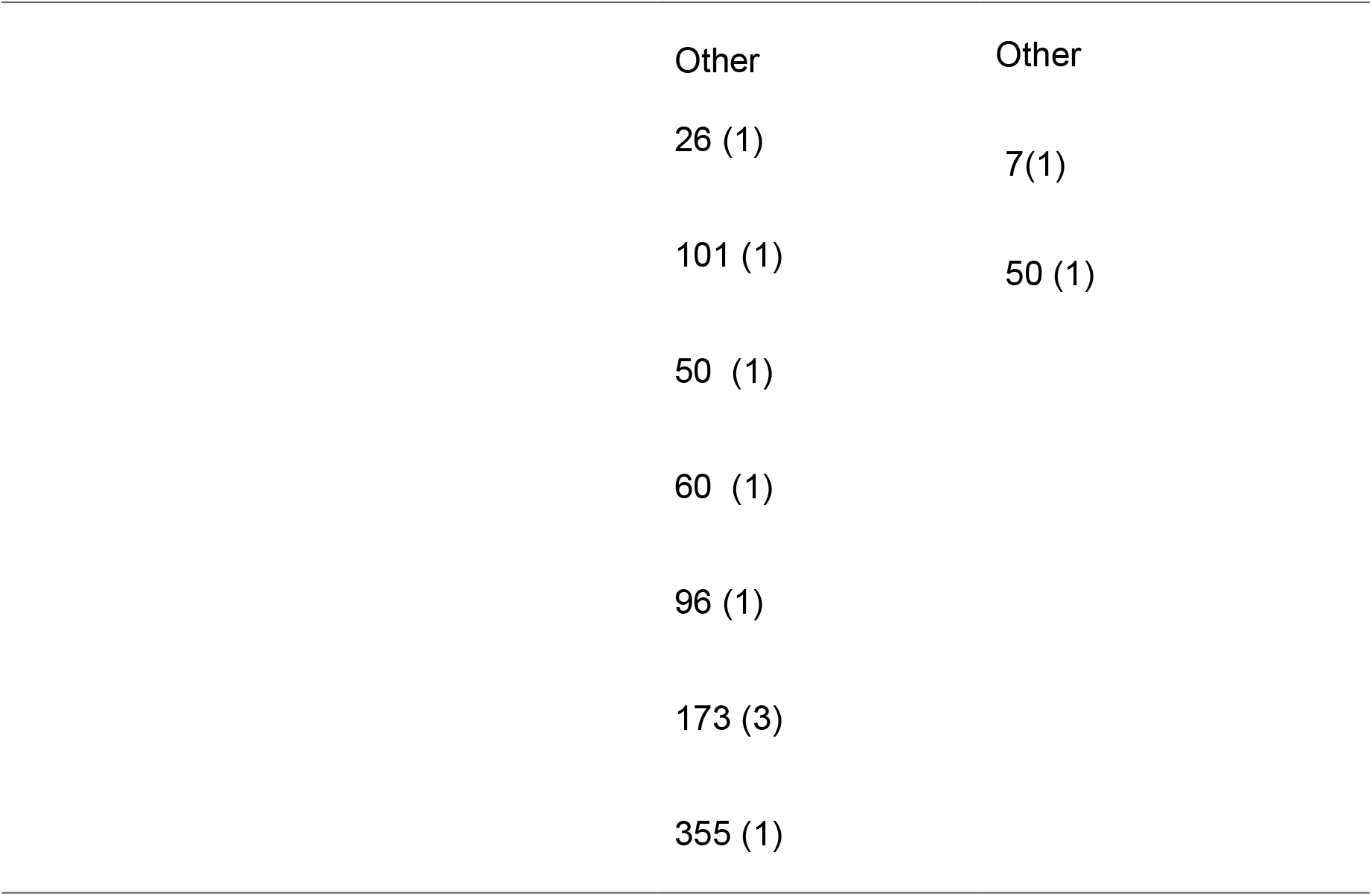
Patient Characteristics.

| **Characteristic** | **Infected** | **Colonized** |
| --- | --- | --- |
| **Number of patients consented, N** | 23 (69.7%) | 10 (30.3%) |
| **Median Age in years** | 70<br>(37 to 86 years) | 66.5<br>(23 to 88 years) |
| **Gender, Male, n** | 13 | 7 |
| **Gender, Female, n** | 10 | 3 |
| **Hospital length of stay (Median), Days (range)** | 20.5<br>(5 to 262 days) | 17<br>(4 to 99 days) |
| **Hospital day blood culture positive for CR-*Kp* (Median), median (range)** | 7.5<br>(0 to 145 days) | 3<br>(0 to 80 days) |
| **Source of Isolate** | Urine (9) | Urine (7) |
|  | Blood (4) | Blood (0) |
|  | Respiratory (5) | Respiratory (2) |
|  | Other | Other |
|  | Abdominal fluid (4) | Rectal Swab (1) |
|  | Penial wound (1) |  |
| **wzi typing** | 29 (7) | 29 (3) |
|  | 154 (7) | 154 (5) |
|  | Other | Other |
|  | 26 (1) | 7(1) |
|  | 101 (1) | 50 (1) |
|  | 50 (1) |  |
|  | 60 (1) |  |
|  | 96 (1) |  |
|  | 173 (3) |  |
|  | 355 (1) |  |

### Anti-CPS antibody responses in clinical patient samples

Patient Ab responses to *wzi29, wzi54* and *wzi50* CPS were measured by ELISA and indicated highest overall IgG responses to CPS *wzi50*, followed by *wzi154*. We also observed minimal IgG response to *wzi29* CPS and that 7 plasma samples lacked any CPS reactivity **(Figure 1A)**. IgM and IgA titers showed similar trends (Supp Fig 2). The magnitude of IgG, IgM, and IgA responses against *wzi29* and *wzi154* were comparable regardless of the state of CR-*Kp* infection (“infected” vs “colonized”). In contrast, higher IgA titers against *wzi50* CPS were only onserved in symptomatic patients **(Supp. Figure 1)**. Plasma (n=33) was also tested for anti-LPS Abs, which have also been observed to be protective (21). Both cohorts exhibited high IgG and low IgA and IgM reactivity against LPS in plasma **(Figure 1B)**.

**Figure 1:**
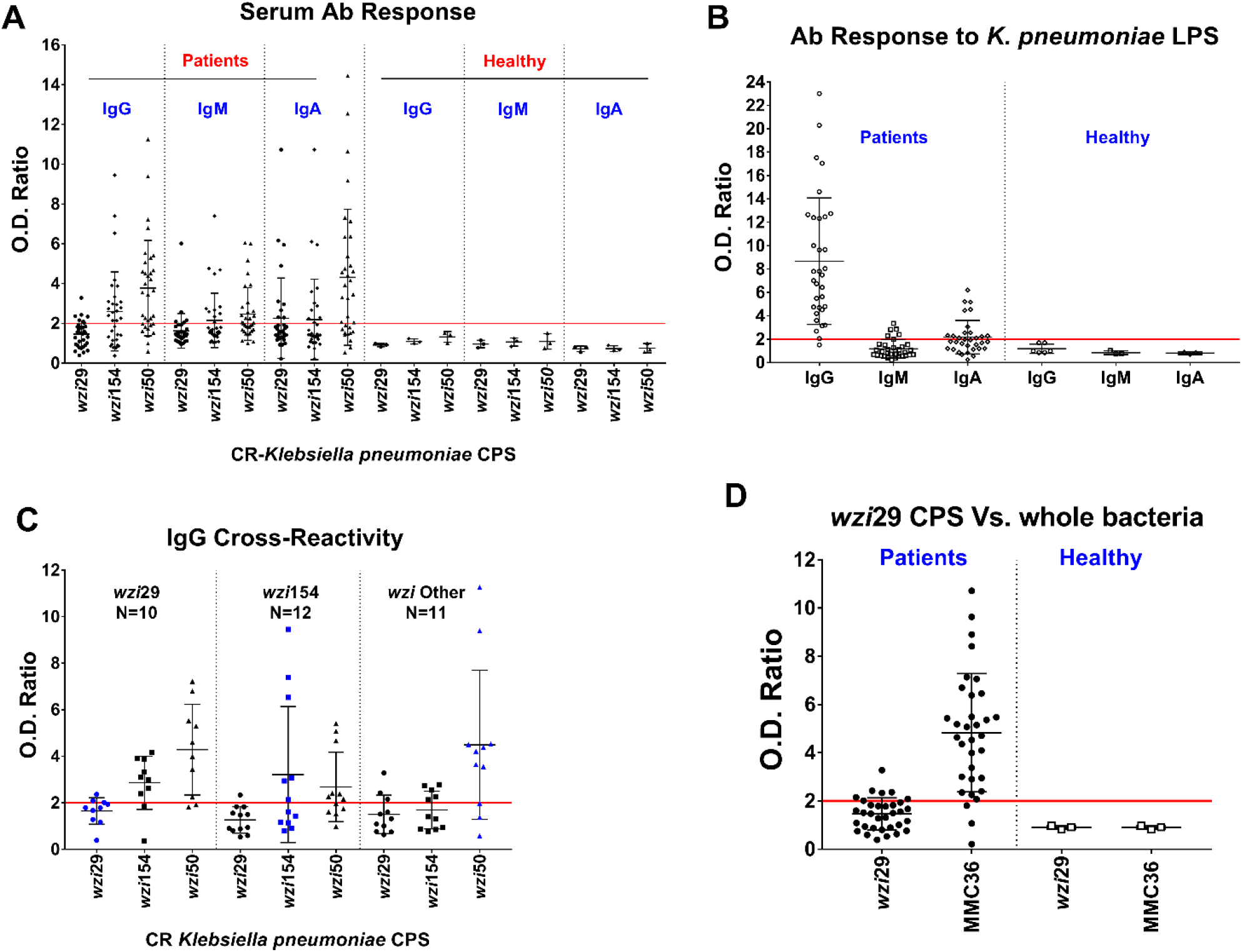
Humoral responses to capsular polysaccharides (CPS) and LPS isolated from carbapenem-resistant *K. pneumoniae*. **(A)** Humoral antibody (IgG, IgM, and IgA) responses detected against *wzi29, wzi154* and *wzi50* CPSs in CR-*Kp* infected patients versus healthy controls. **(B)** Presence of antibodies against *K. pneumoniae* LPS in all CR-*Kp* patients and healthy donors. **(C)** Antibodies of patients infected with *wzi29 K. pneumoniae* (blue dots in the first section) tested against all three CPS types. *wzi154*-infected patients’ antibodies (blue dots in the middle section) tested against all three CPSs. Antibodies from patients infected with other *wzi*-types *CR-K. pneumoniae* (blue dots in the last section) tested against all three CPSs. Blue dots indicate the conditions where the CPS tested and the *wzi* type of the Ab donor’s infection match. **(D)** Presence of antibodies against *wzi29* whole bacteria in CR-*Kp* infected patient plasma Abs versus healthy donors, when compared to purified *wzi29* CPS. Each symbol represents one patient. O.D. Ratio = (O.D. CPS/ O.D. BSA). The fold-change cut-off is set at Y=2 (red line), patients with presence of anti-CPS Abs in the plasma had value ≥2 whereas patients with undetectable level of anti-CPS Abs in plasma had value <2; N = 33; each dot on the scatter plot are individual CR-*Kp* patients’ O.D. ratio and the deviation in O.D. ratios are shown with SD.

Importantly, no Ab reactivity with CPS or LPS was detected in the plasma of healthy individuals, thus suggesting Ab responses were elicited during the recent CR-*Kp* infection **(Figure 1A&B)**. Knowing the *wzi* type of the infecting CR-*Kp* of the patients permitted further subanalysis of Ab reactivity. Notably, no *wzi29*-specific IgG was detected in the plasma of patients infected with *wzi29* CR-*Kp* (**Figure 1C**), although those patients produced cross-reactive Abs that bound to *wzi154* (50%) and *wzi50* (88%) CPS **(Figure 1C)**. While 50% of patients infected with *wzi154* CR-*Kp* mounted a Ab response to *wzi154* CPS, 67% of those patients also exhibited cross-reactive IgG Abs to *wzi50* CPS **(Figure 1C)**. Similarly, while 81% of patients infected with CR-*Kp* strains with other *wzi* CPS (including both *wzi50* infected patients) produced IgGs that recognized *wzi50* CPS, only low Ab titers binding *wzi29* and *wzi154* CPS were observed **(Figure 1C)**. No IgM reactivity to any CPS type was observed in patients infected with *wzi29* CR-*Kp* **(Supp. Figure 2A)**, whereas few patients infected with CR-*Kp* strains with *wzi29* and *wzi154* type CPS had low IgA titers to *wzi29* type CPS **(Supp Figure 2B)**. Lack of any IgG response to *wzi29* type CPS raised the concern that the immunogenic epitopes was destroyed during purification and therefore ELISAs was done with whole bacteria, which confirmed that most patients’ plasma bound to whole *wzi29* MMC36 bacteria **(Figure 1D)**. Isotype subclass analysis showed that anti-CPS Abs included IgG2, IgG3, and IgG4 isotypes, whereas anti-LPS bs were all IgG2 **(Supp. Figure 3)**.

### Cross-agglutination and serum-killing of CR-*Kp* by patient-derived IgGs

To further explore anti-capsular antibody immunity against CR-*Kp*, we examined agglutination and serum resistance of bulk IgG purified from the plasma of a patient who possessed high anti-CPS titers (O.D.> 0.6). First we tested the ability of poly-IgG^#168^ (purified from a patient infected with *wzi50*-producing SBU168) to agglutinate CR-*Kp* strains with different CPS types **(Figure 2 A & B)**. Our data showed that poly-IgG^#168^ agglutinated CR-*Kp* strains MMC34 (*wzi154*), MMC36 (*wzi29*) and MMC38 (*wzi50*) **(Figure 2A)**. The extent of agglutination (average area of agglutinated clumps) was quantified by ImageJ software and was found to be significant for strains MMC34 and MMC38 when compared to bacteria treated with IgG derived from normal human serum (poly-IgG^NHS^) (unpaired t-test, P=0.0040) **(Figure 2B)**. Next, serum resistance or killing was evaluated over a 3-hour incubation time. CR-*Kp* strains differ in their resistance to complement-mediated killing, whereby some strains can be killed whereas for other strains only growth is inhibited (8). For the complement-sensitive strain MMC34, poly-IgG^#168^ induced killing and significantly (P<0.0001) inhibited bacterial growth in 75% normal human serum when compared to NHS and poly-IgG^NHS^ controls **(Figure 2C)**. Our data indicate that co-incubation of poly-IgG^#168^ significantly impaired growth of MMC36 in 75% NHS **(Figure 2D)** compared to serum with non-specific poly-IgG^NHS^ (P<0.0001). MMC38 treated with poly-IgG^NHS^ had an overall 145% growth increase, whereas negligible growth was observed in MMC38 treated with poly-IgG^#168^ between the time interval of 2 and 3 hours (P<0.0001) **(Figure 2E)**

**Figure 2:**
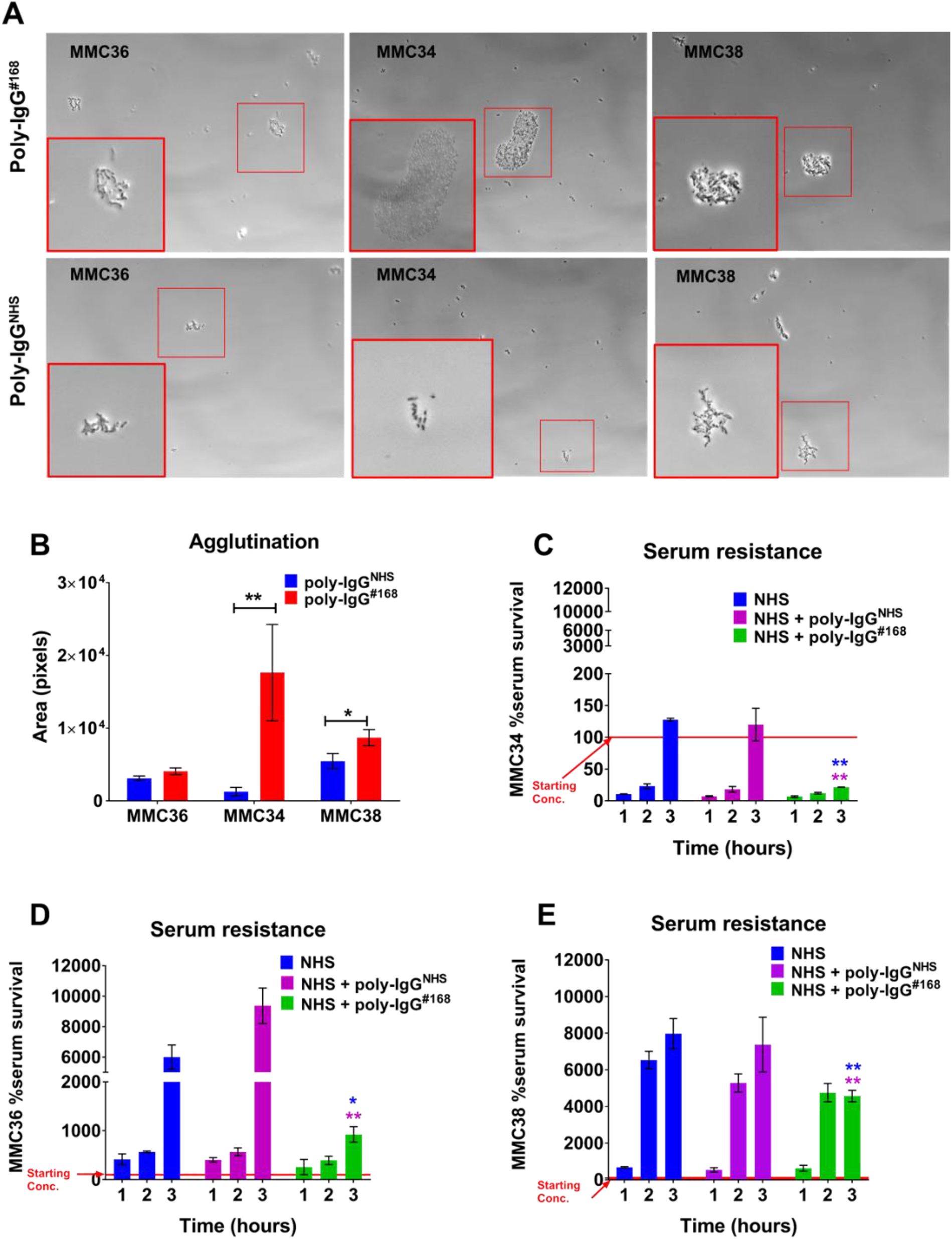
Patient antibodies promote agglutination and decrease serum resistance. **(A)** *wzi*50 patient-derived (PD) IgGs promoted agglutination of *wzi*29 MMC36, *wzi*154 MMC34, and *wzi50* MMC38 strains, which was visualized with phase-contrast microscopy at 200X. **(B)** Quantification of the area of agglutinated bacteria. **(C)** *wzi50* PD IgGs (poly-IgG^#168^) mediated serum killing of *wzi*154 MMC34; **(D)** inhibited the growth of *wzi29* MMC36; and **(E)** inhibited the growth of *wzi50* MMC38 strains, with-respect-to CR-*Kp* strains treated with poly-IgG^NHS^ in 75% NHS. Bars depict means and SD of three independent experiments. Overall differences between treatment groups were determined to be significant by Repeated-Measures Two Way ANOVA using Tukey’s post-hoc test displayed in-graph. Bacterial growth in serum cut-off normalized to Time=0 hour is set at Y=100 (red line). For all in-graph statistics, p-values displayed in blue are comparisons to the NHS only control group, whereas p-values in purple are comparison to the poly-IgG^NHS^ treated group. p values are replaced with ns if >0.1 (ns); * if < 0.05; ** if <0.01; and *** if p<0.001.

### Opsonophagocytosis of clinical CR-*Kp* isolates by patient-derived poly-IgG

A key attribute commonly associated with efficacy of CPS specific Ab-mediated immunity is opsonophagocytosis (22, 23). To assess the efficacy of different PD poly-IgGs, we performed opsonophagocytosis experiments in J774 macrophages with Poly-IgG^#207^, poly-IgG^#219^, and poly-IgG^#116^ purified from patients infected with SBU219 (*wzi154*), SBU116 (*wzi50*), and SBU207 (*wz*i*29*). Opsonophagocytosis of respective CR-*Kp* clinical isolates was promoted in a dose-dependent matter by all 4 poly-IgGs **(Fig 3A-C and sup Fig 4)**. Due to limited quantity of poly-IgG^#116^, investigations were continued with poly-IgG^#116^ also derived from a patient infected with *wzi50* producing CR-*Kp*. Poly-IgG^#207^ promoted opsonophagocytosis of two different *wzi29* strains SBU207 and MMC36 **(Figure 3D)**. In a similar fashion poly-IgG^#219^ **(Figure 3E)**, and poly-IgG^#116^ **(Figure 3F),** induced opsonophagocytic uptake of MMC34 and SBU219 (both *wzi154*:); and MMC38 and SBU116 (both *wzi50*) strains respectively. Interestingly, while poly-IgGs promoted phagocytosis of capsular strains including 33576 (*wzi154*) it did not mediate phagocytosis of its acapsular 33576 Δ*wzy* mutant **(Figure 3 D-F)**. These data suggest that anti-CPS Abs in PD poly-IgGs convey opsonophagocytic efficacy and also indicate presence of cross-reactive anti-CPS Abs.

**Figure 3:**
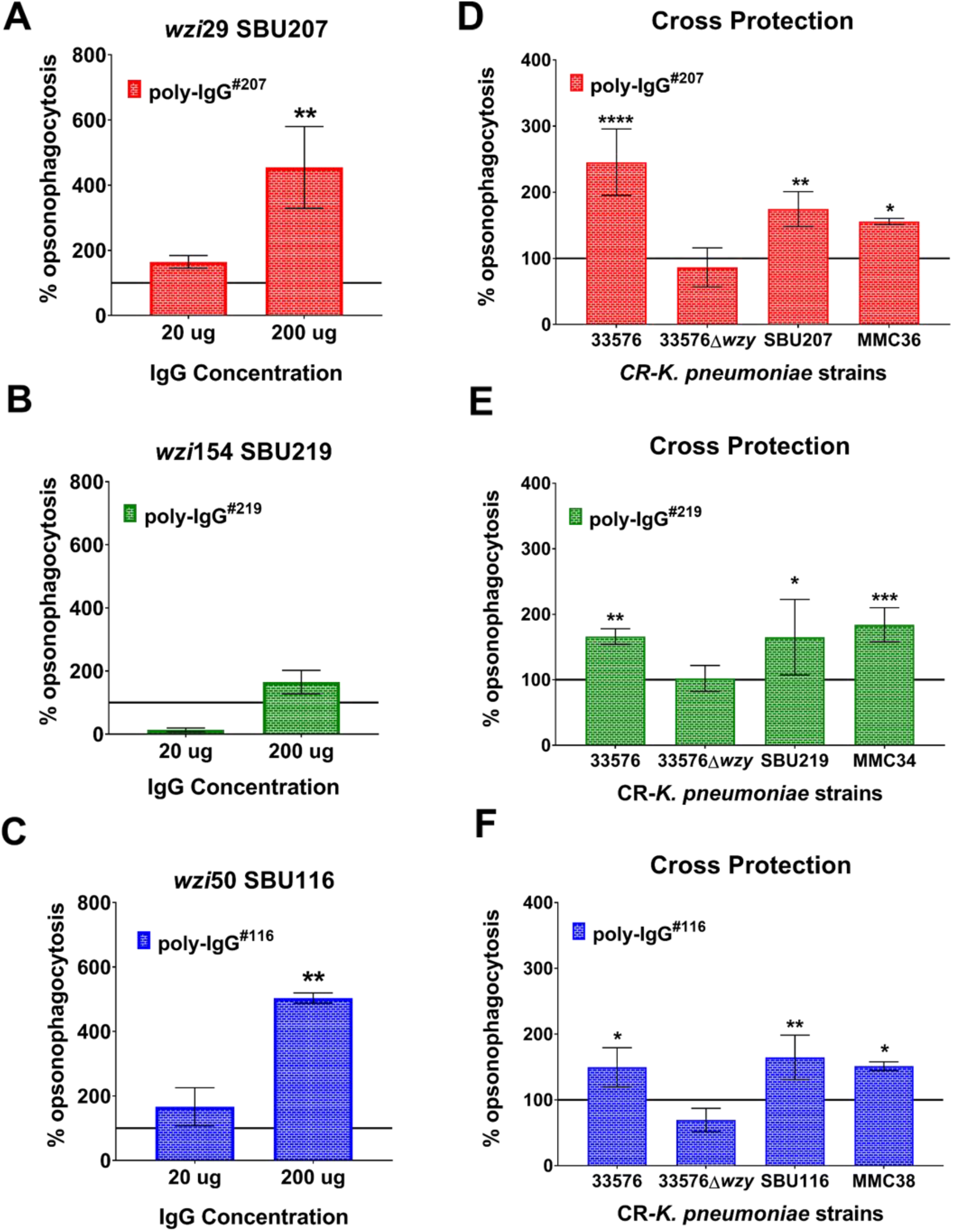
Coincubation with patient derived IgGs induce opsonophagocytosis of clinical *K. pneumoniae* strains. PD IgGs isolated from patient #207 (*wzi29*), 219 (*wzi154*), and #116 (*wzi50*) induced opsonophagocytosis of patient-matched *K. pneumoniae* **(A)** SBU207 (*wzi29*), **(B)** SBU219 (*wzi154*), and **(C)** SBU116 (*wzi50*) strains, respectively. PD IgGs **(D)** poly-IgG^#207^, **(E)** poly-IgG^#219^, and **(F)** poly-IgG^#116^ broadly recognized and induced opsonophagocytosis of strains with similar *wzi* type and also cross-reacted with 33576 clade2 *Kp* capsular strain but did not mediate phagocytosis of acapsular 33576Δ*wzy* mutant strain. Bars depict means and SD of three independent experiments, with wells performed in triplicate. Phagocytosis cut-off normalized to PBS control is set at Y=100 (**black line**), positive phagocytosis had value >100 and no phagocytosis or inhibition of phagocytosis ≤ 100. Overall differences between treatment groups were determined to be significant by Repeated-Measures Two Way ANOVA using Tukey’s post-hoc test displayed in-graph. For all in-graph statistics, p values displayed in black are comparisons to the phagocytosis of acapsular 33576Δ*wzy* mutant strain. p values are replaced with ns if >0.1 (ns); * if < 0.05; ** if <0.01; and *** if p<0.001.

### Relevance of anti-capsular antibodies for opsonophagocytic efficacy

To test if the protective Abs are anti-CPS we depleted poly-IgG^#116^ of *wzi154* and *wzi50* CPS-specific Abs by co-incubation with corresponding CPS-coated beads. Ab-mediated opsonophagocytosis of 4 CR-*Kp* strains producing distinct *wzi* types (*wzi29, wzi154*, and *wzi50*) was compared to that of non-depleted poly-IgG^#116^ and poly-IgG^#NHS^. Data confirmed significant augmented phagocytosis of all 4 CR-*Kp*s **(Figure 4 A-D)** when compared to poly-IgG#^NHS^. For MMC34, MMC8 and SBU116 phagocytosis was significantly lower when PD bulk IgGs was depleted with either *wzi154*- or *wzi50*- CPS-coated beads **(Fig 4 A-C**). Depletion of CPS-specific Abs from poly-IgG^#116^ had no effect on opsonophagocytosis of *wzi29* MMC36 strain (**Figure 4D**). Similar results were observed when *wzi154* specific Abs were depleted from bulk poly-IgG^#219^ resulting in 40% and 30% loss in phagocytic efficacy for MMC34 and MMC38 respectively **(Figure 4 E and F)**, whereas depletion of *wzi50* specific Abs only affected the phagocytosis of MMC34 (36% loss) **(Figure E)**. Depletion of either anti-CPS Abs did not affect the phagocytosis of MMC36 (data not shown).

**Figure 4:**
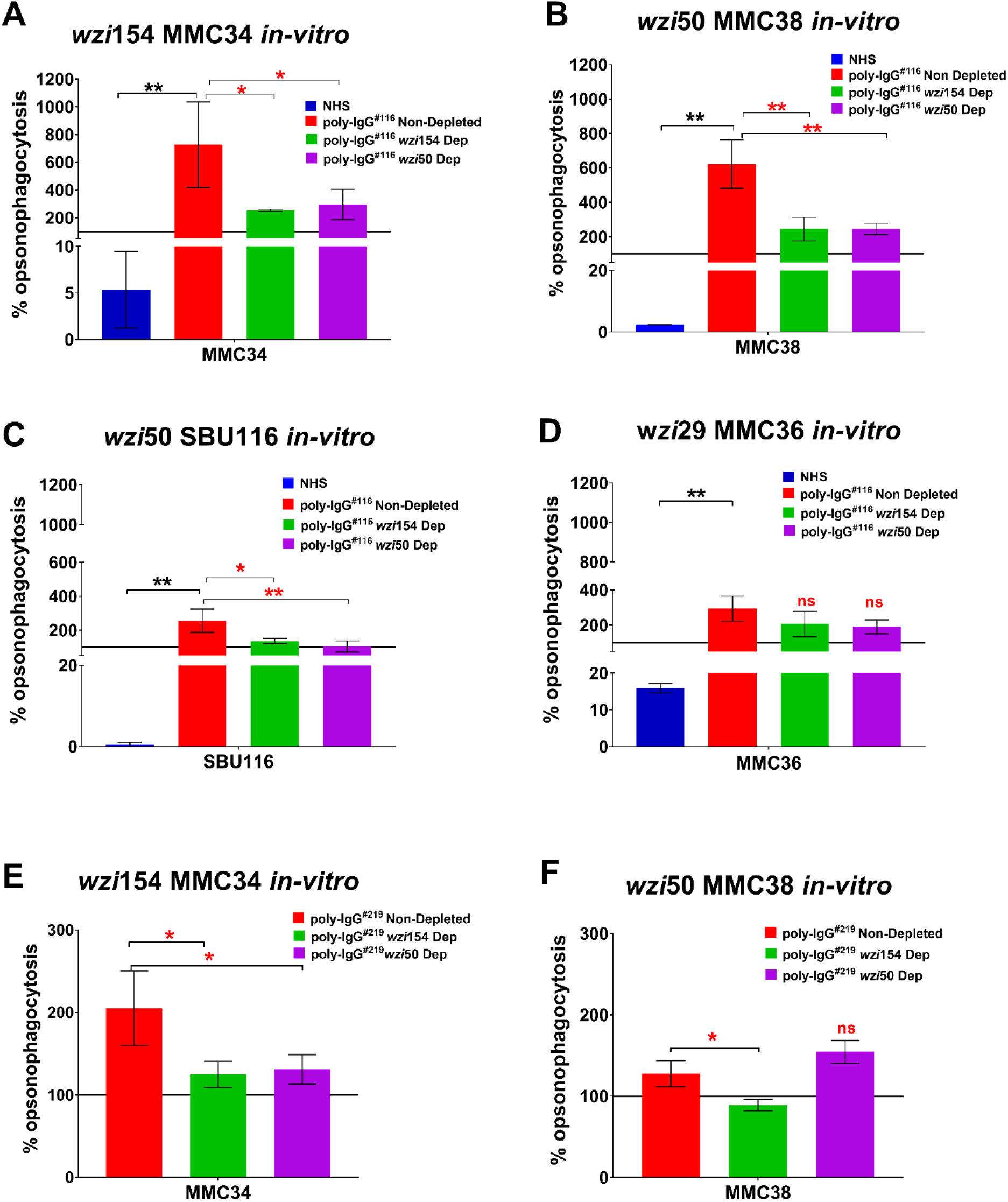
Depletion of CPS-specific antibodies inhibits oposonophagocytosis of corresponding *K. pneumoniae*. PD-IgGs poly-IgG^#116^ and poly-IgG^#219^ provided cross-protection against all clade types (including clade1 & clade2). Depletion of *wzi154* and *wzi50* CPS specific IgGs from poly-IgG^#116^ significantly inhibited opsonophagocytosis of **(A)** *wzi154* MMC34, **(B)** corresponding *wzi50* MMC38 and **(C)** patient matched SBU116 strains, but **(D)** effect on opsonophagocytosis of *wzi29* MMC36 was negligible. Depletion of *wzi154* and *wzi50* CPS specific IgGs from poly-IgG^#219^ significantly inhibited opsonophagocytosis of **(E)** corresponding *wzi154* MMC34 strain, whereas **(F)** depletion of *wzi154* CPS only inhibited the opsonophagocytosis of *wzi50* MMC38. Bars depict means and SD of three independent experiments, with wells performed in triplicate. Phagocytosis cut-off normalized to PBS control is set at Y=100 (black line), positive phagocytosis had value >100 and no phagocytosis or inhibition of phagocytosis ≤ 100. Overall differences between treatment groups were determined to be significant by Repeated-Measures Two Way ANOVA using Tukey’s post-hoc test displayed in-graph. For all in-graph statistics, p values displayed in black are comparisons to the NHS group, whereas p values in red compare poly-IgGs Non-depleted with *wzi154* and *wzi50* depleted Abs. p values are replaced with ns if >0.1 (ns); * if < 0.05; ** if <0.01; and *** if p<0.001.

### Reduced protective efficacy of depleted PD-poly IgG in a murine infection model

Lastly, we explored if cross-reactive Abs could protect mice from CR-*Kp* infection. We compared CFU in mice pulmonary model infected with different CR-*Kp* strains (MMC34, SBU116, and MMC36) either pre-opsonized with undepleted or depleted poly-IgG^#116^. Mice injected with poly-IgG^#116^-opsonized MMC34 exhibited moderate 1-log_10_ reduction in bacterial lung burden as well as 4-log_10_ fold reduction in dissemination to spleen. **(Figure 5)**. A 2-log_10_ and 1-log_10_ reduction in lung and spleen CFU respectively was seen in mice infected with poly-IgG^#116^ -opsonized SBU116. Depletion of *wzi50*-specific Abs resulted in 1-log_10_ higher CFU in both lung and spleen when compared to undepeted treatment. Whereas *wzi154* specific Ab depletion only affected dissemination to spleen when compared to undepleted treatment **(Figure 5)**. A significant albeit only small decrease in lung CFU was observed in mice infected with MMC36 pre-opsonized with poly-IgG^#116^, but no dissemination to spleen was observed in either experimental groups. Furthermore, bacterial burden in lung was identical between *wzi154* depleted group and PBS control **(Figure 5)**.

**Figure 5:**
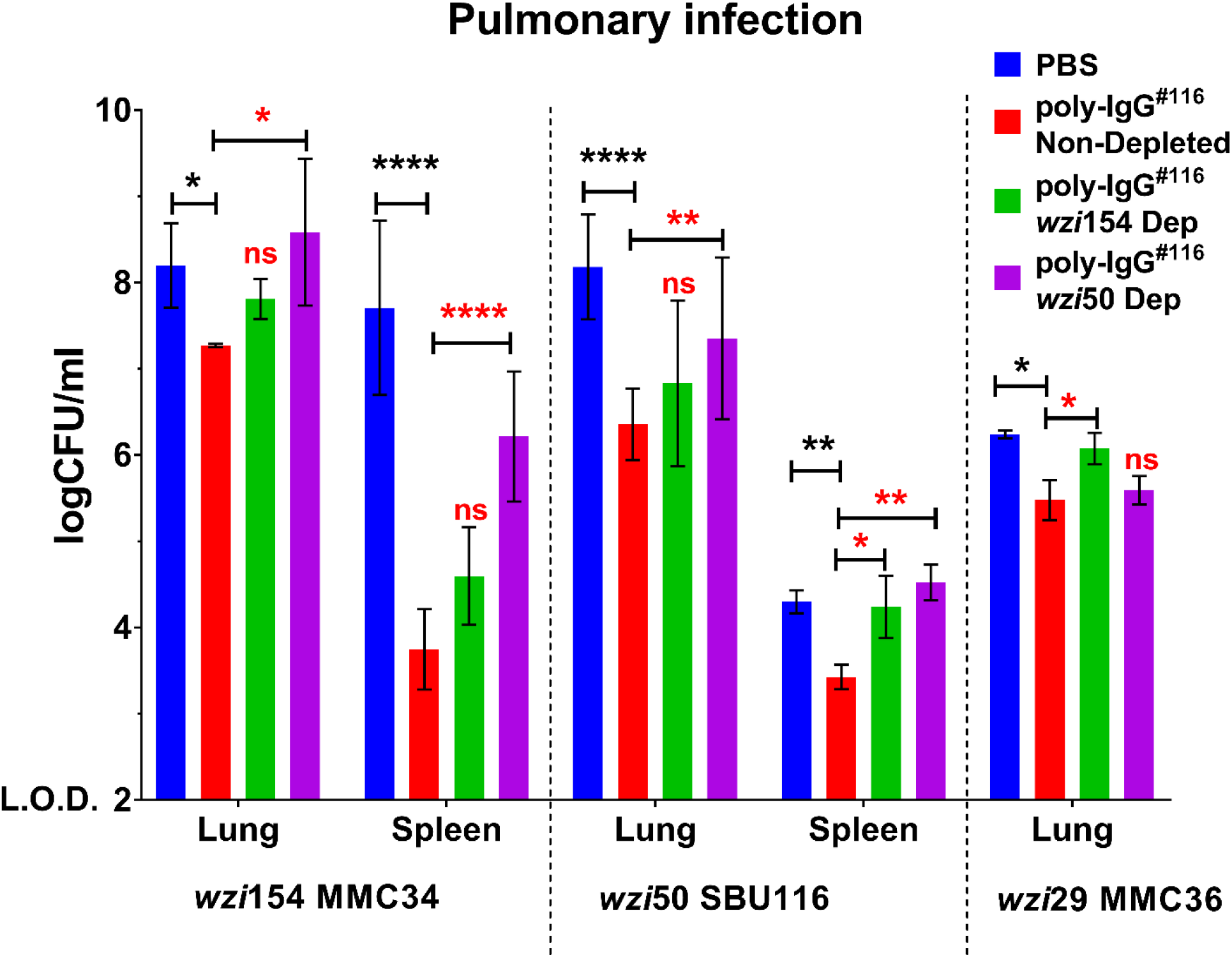
Passive transfer of purified human polyclonal IgG from CR-*Kp* subjects reduces K. pneumoniae bacterial burden in CR-*Kp*-infected mice, whereas specific-CPS depletion reverses the therapeutic effect. Bacterial burden in lungs, and spleen of mice infected with a lethal inoculum of MMC34, SBU116 and MMC36 strains pre-opsonized with either CPS-specific depleted or non-depleted PD-IgGs. For all studies, bars depict means and SD for overall differences in CFU between treatment groups (n=3 per group) were assessed for significance by Two Way-ANOVA with multiple-comparison correction and limit of detection (L.O.D) was set at Y=2. For all in-graph statistics, p values displayed in black are comparisons to the PBS group, whereas p values in red compare poly-IgGs Non-depleted with *wzi154* and *wzi50* depleted Abs. p values are replaced with ns if >0.1 (ns); * if < 0.05; ** if <0.01; and *** if p<0.001.

## Discussion

Prior studies in mice and monkeys provide compelling evidence that CPS-specific Abs are effective against CR-*Kp* (15, 16, 19, 24). Several groups have developed vaccines to prevent *K. pneumoniae* infection that target CPS (16, 25–27). However, concerns regarding feasibility prevail because of the heterogeneous polysaccharide capsule (8). These data constitute the first analysis of the anti-CPS Ab response in CR-*Kp* infected patients. Several important conclusions can be drawn from our data. First, most CR-*Kp* infected patients mount a humoral response to the polysaccharide capsule, albeit it is more variable in its magnitude than Ab reactivity to LPS. Second, CPS-specific Abs cross-react with other capsule types. Third, protective Abs were specific to CPS. Lastly, poly-IgG from CR-*Kp* infected patients can protect CR-*Kp* infected mice, and loss of efficacy is observed after depletion of CPS-specific Abs.

Our data represents an unbiased analysis of the Ab response in CR-*Kp* infected patients as the cohort included the majority of hospitalized patients diagnosed with CR-*Kp* infection during the respective time period. Although colonized patients were diagnosed earlier and spent less time in the hospital, there were no striking differences with-respect-to their CPS-specific Ab response. We chose to further analyze the Ab response to the three most common *wzi* types of ST258 strains. The *wzi154* CPS type is expressed by most clade 2 strains, *wzi29* is the most prevalent CPS expressed by clade 1 strains and *wzi50* can be expressed by clade 1 as well as clade 2 strains (8, 9, 28). Interestingly, three of the seven patients who did not mount a CPS-specific Ab response were infected with less common *wzi*-types (*wzi173* and *wzi7*).

Despite the CPS variability in CR-*Kp*, many of the patients’ plasma reacted to more than one CPS. This was not necessarily expected because the structure of *wzi154* (MMC34) is distinct from other published *K. pneumoniae* CPS structures (29) and its sugar composition is also very different from that of *wzi50* (8). Regardless, about half of the patients infected with *wzi154* CR-*Kp* strains mounted high titers to *wzi154* and *wzi50* CPS. Interestingly, plasma derived from patients infected with *wzi29* CR-*Kp* exhibited the most pronounced cross-reactivity with other *wzi*-types but no reactivity with *wzi29* CPS. Although this result was consistent with published studies reporting failure of *wzi29* CPS to elicit Abs in mice or rabbits (16, 17), it was not in accord with results from *in vitro* phagocytosis assays. They indicated opsonophagocytic efficacy of PD-poly-IgG of *wzi29* CR-*Kp* strains, whereas acapsular mutants were not phagocytosed, consistent with the presence of Abs specific for *wzi29* type CPS. Although these results are not conclusive, a modified ELISA demonstrating Abs to whole *wzi29* CR-*Kp* further support the conclusion that *wzi29* type CPS indeed elicits Abs. We hypothesize that critical CPS epitopes are destroyed during CPS purification and better methods to purify CPS are needed. Cross-reactive Abs have been identified in the serum of volunteers vaccinated with CPSs derived from different *K. pneumoniae* (30). Such cross-reactivity may be the result of cocktails of distinct Abs that bind to different antigens or alternatively some Abs could react with more than one CPS. Indeed, one study, reported that 12% of human mAbs cloned from subjects vaccinated with 23-valent Pneumovax cross-reacted with two serotypes (31). Although our murine mAbs exclusively bind to *wzi154* type CPS, one could potentially clone hybridomas that produce cross reactive Abs through modified vaccination protocols and altered screening procedures,.

Importantly, healthy subjects did not exhibit Ab reactivity with *Kp*-derived CPS or LPS, indicating that these Ab reactivities are the result of infection and colonization with CR-*Kp*. Further studies will need to resolve if CPS-specific IgA is present in the colon. Interestingly, the dominance of a specific IgG subclass was not observed for CPS-specific Abs whereas the LPS of *Klebsiella* elicited predominantly IgG2 response. Although one vaccine study in mice with CR-*Kp* oligosaccharides identified the murine subclass mIgG1 to be the most seroprevalent in those mice (32), human IgG2, similar to the analogous murine isotype (mIgG3), is the primary humoral response to T-independent antigens such as CPS. This IgG subclass preference was also observed in humans infected with *Helicobacter pylori* and *Mycobacterium tuberculosis* (33, 34).

Abs directed against the O polysaccharide are protective against wild-type *Kp* in lethal infection models (35, 36). Abs that bind to LPS have not only been shown to be opsonophagocytic but also cross-reactive (21). Experiments with acapsular mutants as well as studies with bulk plasma that was depleted of CPS-specific Abs were, therefore, necessary to show the causal role of anti-CPS antibodies in opsonophagocytic efficacy. The first set of experiments demonstrate that bulk IgG promotes agglutination of two CR-*Kp* strains (*wzi50, wzi154*) and enhances serum resistance in all three *Klebsiella* strains, which differ in agglutination and serum resistance (13, 16, 17). Ab-mediated complement deposition of C5b-9 and C3c has been shown to be critical for Ab activity (16). Enhancement of opsonophagocytic efficacy by two different high-titer IgG bulk fractions demonstrated that enhancement of phagocytic activity was mediated by CPS specific Abs and lost after depletion. Again, cross-reactivity was observed for MMC34 (*wzi154*) as both depletion of *wzi50*, as well as *wzi154* specific Abs, affected the phagocytic efficacy.

Regrettably, CR-*Kp* strains exhibit low virulence and *in vivo* experiments continue to be a challenge in mice (16, 37) and even cynomolgus macaques (19). *In vivo* efficacy of poly-IgG^#116^ a pulmonary infection model further support our conclusion that infected patients generate a protective humoral immune response and that CPS-specific Abs are relevant Abs in both bulk IgG fraction. PD-Poly-IgG^#116^ lowered bacterial burden in the lung and dissemination to the spleen in mice infected with MMC34, MMC36, and MMC38, which expressed *wzi154, −29* and −*50* type CPS, respectively. Importantly, protective efficacy was reversed when CPS-specific Abs were depleted underscoring their relevance.

In summary, despite marked capsular heterogeneity, our results demonstrate that infection with CR-*Kp* induces cross-reactive anti-CPS Abs. The finding that most (79%) elderly patients irrespective of their state of infection mounted an Ab response to at least one of the three CPS underscores the immunogenicity of CPS and indicates feasibility of a CPS-based vaccine strategy that targets the elderly who are also the most vulnerable. In contrast to *S. pneumoniae*, CR-*Kp* colonization is not yet widely spread in the community and the majority of patients still become colonized through nosocomial exposure in diverse health care settings (38, 39). Recent advances with recombinant production of bioconjugate vaccines in glycoengineered *Escherichia coli* cells, against the 2 *Kp* serotypes, K1 and K2 could improve generation of multivalent CPS-based vaccines (24). Our data indicate that vaccines with *wzi154, wzi29*, and *wzi50* could potentially give broad coverage which is preferablebecause similar to pneumococcal vaccines non-vaccine CPS-types could increased after the introduction (40, 41). In summary, our results encourage efforts to further develop CPS-based vaccines. Furthermore, these data suggest that certain CPS may elicit more cross-reactive Abs but care has to be taken when purifying CPS to conserve immunogenic epitopes.

## Materials and Methods

### Ethics statement

Animal study protocols were approved by the Animal Committee (IACUC) at SBU (approval no. 628253). This study is in strict accordance with federal, state, local, and institutional guidelines that include the Guide for the Care and Use of Laboratory Animals, the Animal Welfare Act, and Public Health Service Policy on Human Care and Use of Laboratory Animals. All surgery was performed under ketamine-and-xylazine anesthesia, and every effort was made to minimize suffering. Patients were consented under institutional review board (IRB) and SBU Human Subjects Committee approved protocols (IRB# 896845 and 851803). Healthy donors gave written informed consent for blood donation under IRB# 718744.

### Collection of plasma and CR-*Kp* from patients

Patients admitted to Stony Brook University Hospital (SBUH) within 2017-2019, from whom CR-*Kp* was isolated, were consented and their CR-*Kp* strains and plasma were collected and frozen at −80°C and −20°C, respectively. Patients were classified as being “colonized” or “infected” based on standard criteria described elsewhere (12).

### CR-*Kp* strain isolation, *wzi* typing, and purification of capsular polysaccharide

CR-*Kp* strains were identified by standardized methods according the Clinical and Laboratory Standards Institute (CLSI) as well as revised CDC criteria (4) and *wzi*-typed according to published protocols(6). For experiments, CR-*Kp* strains isolated from patients as well as MMC36 (*wzi29*), MMC34 (*wzi154*) and MMC38 (*wzi50*) (8), were cultured in Luria-Bertani (LB) broth and agar at 37°C (8). The CPS of the previously characterized three strains was purified for ELISA and depletion studies based on a previously-described protocol (15). Absence of lipopolysaccharide (LPS) (< 20EU/ml (42)) in purified CPS was confirmed by Pierce™ LAL Chromogenic Endotoxin Quantitation Kit (Thermo Scientific).

### Detection of anti-CPS and anti-LPS antibodies in plasma by CPS or LPS ELISA

ELISA-assays described in (15) were employed to detect circulating CPS Abs in the plasma of patients. Plasma samples were added in 1:50 dilution (determined by titration 1:25 to 1:150) for all ELISAs and secondary anti-human IgG, IgM, IgA and IgG subclass antibodies were used for detection. Presence of CPS-specific Abs was defined by O.D. ratio value ≥ 2 (O.D ratio = O.D._405_ CPS/ O.D._405_ BSA) where an O.D. ratio value < 2 was considered non-specific. For detection of lipopolysaccharide (LPS) specific Abs, a similar ELISA assay was done using *K. pneumoniae* LPS purchased from Sigma-Aldrich (L4268).

### Bulk IgG isolation

Bulk patient-derived (PD) poly-IgG (PD IgGs) was purified from plasma of patients infected with SBU116, SBU168, SBU207, and SBU219 respectively or normal human serum (NHS) (H4522, Millipore Sigma) by negative selection using Melon Gel resin (Thermo Scientific) according to manufacturer’s protocol. Bulk IgGs were quantified using Human IgG ELISA development kit (Mabtech).

### Bulk IgG serum resistance assay and Rapid agglutination assay

Human serum resistance assay was done with poly-IgG^#168^ (40 μg/ml) by modifying the protocol described in (43). Agglutination assays of MMC34, MMC36, and MMC38 strains was carried out with NHS-derived IgG (poly-IgG^NHS^) or with PD IgGs isolated from a patient infected with SBU168 (40 μg/ml) as described (15, 16). Briefly 3 × 10^6^ CR-*Kp* strains were incubated with poly-IgG^#168^ in 75% NHS at 37°C for 0, 1, 2- & 3-hour time intervals. At each interval, samples were diluted serially and plated on LB plates. The experiment was repeated thrice.

Agglutination assays of MMC34, MMC36, and MMC38 strains was carried out with NHS-derived IgG (poly-IgG^NHS^) or with PD IgGs isolated from a patient infected with SBU168 (40 μg/ml) as described (15, 16). Agglutination was captured in phase-contrast at a magnification of 200X with Zeiss deconvolution microscope and images were analyzed with ImageJ software. The experiment was repeated thrice.

### Whole-Cell ELISA

A whole-bacterium ELISA-assay was used (24) to detect Abs to *wzi*-29-type CPS expressing MMC36. Briefly, half-of the 96-well plates (48-wells) were coated with 8×10^8^ CFU mL^−1^ of *wzi*-29 MMC36 strain in methanol for 24 hours. After 24 hours, the plate was blocked with 2% BSA and a standard anti-CPS ELISA was performed. Presence of Abs in plasma was defined as a O.D. ratio value ≥ 2 and plasma with O.D. ratio value < 2 were considered to have no/negligible anti-MMC36 antibodies

### Biotinylation of capsular polysaccharides and depletion of CPS-specific IgGs

CPS biotinylation was done by adapting the protocol described previously (44). Briefly, purified CPS was constituted in endotoxin-free water at 1mg/ml concentration. 200μl of 100mM sodium periodate was added per 1mg CPS and the solution was incubated for 6 hours. Then, 100μl of 25% glycerol was added to react with excess sodium periodate, and the solution was left at room temperature in the dark for 2 hours. After periodate clean-up, the reaction mixture was passed through 2 Bio-Gel P-2 mini columns (1504114, Bio-rad) by centrifugation (3000 rpm, 2mins) as per manufactures instructions. Biotin-Hydrazide (SP-1100, Vector Laboratories) was dissolved in DMSO at 50mg/ml. For labeling 40μl (2mg) of this solution was mixed with 1mM of manganese dichloride per 1ml of CPS solution and incubated overnight at room temperature. Modified polysaccharides were then passed through two mini columns of Bio-gel P2 and reduced with 1mM sodium borohydride for 5 min at room temperature. The reduced polysaccharide was passed through two Bio-gel P2 mini columns and the level of biotinylation was detected with Pierce Biotin Quantitation Kit (28055, Thermo Scientific).

200μl of Pierce™ NeutrAvidin™ Agarose beads (29200, Thermo Fisher Scientific) were coated with 1mg/ml Biotinylated CPS at 4°C for 16 hours. Coated beads were washed with 2 column volumes of PBS (pH 7.4) and then incubated with bulk PD IgGs at 4°C for 16 hours. Beads were washed with 2 column length volume (2ml) of PBS (Corning) and the CPS-depleted pools were eluted with PBS for further use. Depletion was confirmed by CPS ELISA before use in further experiments.

### Macrophage phagocytosis

Macrophage cell line J774A.1 (ATCC) was used for macrophage phagocytosis assays with bulk PD poly-IgGs (40 μg/ml) using published protocols (15, 16). poly-IgG^#168^, poly-IgG^#116^, poly-IgG^#207^, and poly-IgG^#219^ were used to study antibody-mediated opsonophagocytosis of CR-*Kp* strains (SBU168, SBU116, SBU207, SBU219, MMC34, MMC36, and MMC38); and of *wzi154* CPS-expressing 33576 and acapsular 33576 Δ*wzy* mutant (1). Phagocytosis assay was additionally carried out with CPS-specific depleted PD IgGs (*wzi50* and *wzi154* depleted poly-IgGs) and poly-IgG^#NHS^ (negative control).

### Pulmonary infection model in mice

Pulmonary infection model study was done as per the protocol described in (15). Female C57BL/6 mice 6 to 8 weeks old were used. 2 × 10^8^ CFU/ml CR-*Kp* strains (MMC36, MMC34 and SBU116) were incubated with PD poly-IgGs, CPS-specific-depleted PD poly-IgGs, or PBS at 5 mg/ml for 1 h. A 50μl volume of this inoculum, containing 10^7^ CFU was injected intra-tracheally. After 24 h, mice were euthanized, and lungs and spleens were processed for enumeration of bacteria in homogenized tissue, and bacterial dissemination analysis.

### Statistical analysis

Statistical tests were performed with GraphPad Prism 6 for Windows. For multigroup comparisons of parametric data (e.g., phagocytosis, and serum-resistance assay), ANOVA with post-hoc analysis using Tukey’s, Sidak’s, or Dunnett’s comparison test was used. For two-group comparisons of parametric data, paired t-tests corrected for multiple comparisons using the Holm-Sidak method were performed.

## Acknowledgment

We would like to thank Liang Chen and Barry Kreiswirth for gifting us CR-*Kp* strains 33576 and 33576 Δ*wzy* mutant carbapenem-susceptible ST258 *K. pneumoniae* strains (1). Additionally, we would like to thank Somanon Bhattacharya for his assistance in editing the manuscript.

## SUPPLEMENTARY FIGURES LEGENDS

**Supplementary Figure 1:**
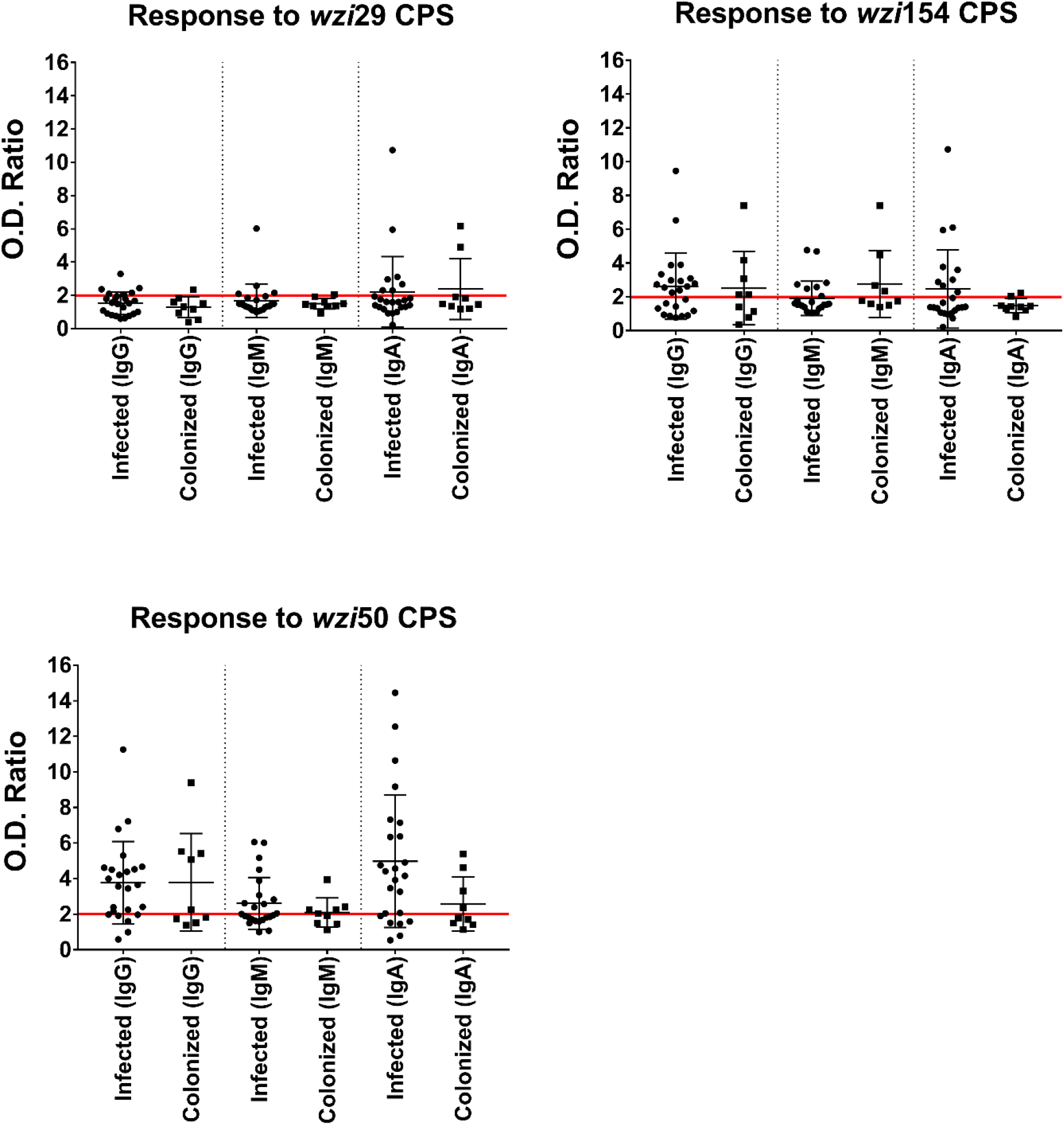
Antibody responses in “infected” versus “colonized” CR-*Kp* patients to capsular polysaccharides (CPS) isolated from CR-*Kp* strains. Humoral responses to CPS were independent of the patient’s clinical status. O.D. Ratio = (O.D. CPS/O.D. BSA). The positive cut-off is set at Y=2 (red line), the positive serum had the value ≥2 and negative serum is <2; N=33; each dot on the scatter plot are individual CR-*Kp* patients’ O.D. ratio and the deviation in O.D. ratios are shown with SD.

**Supplementary Figure 2:**
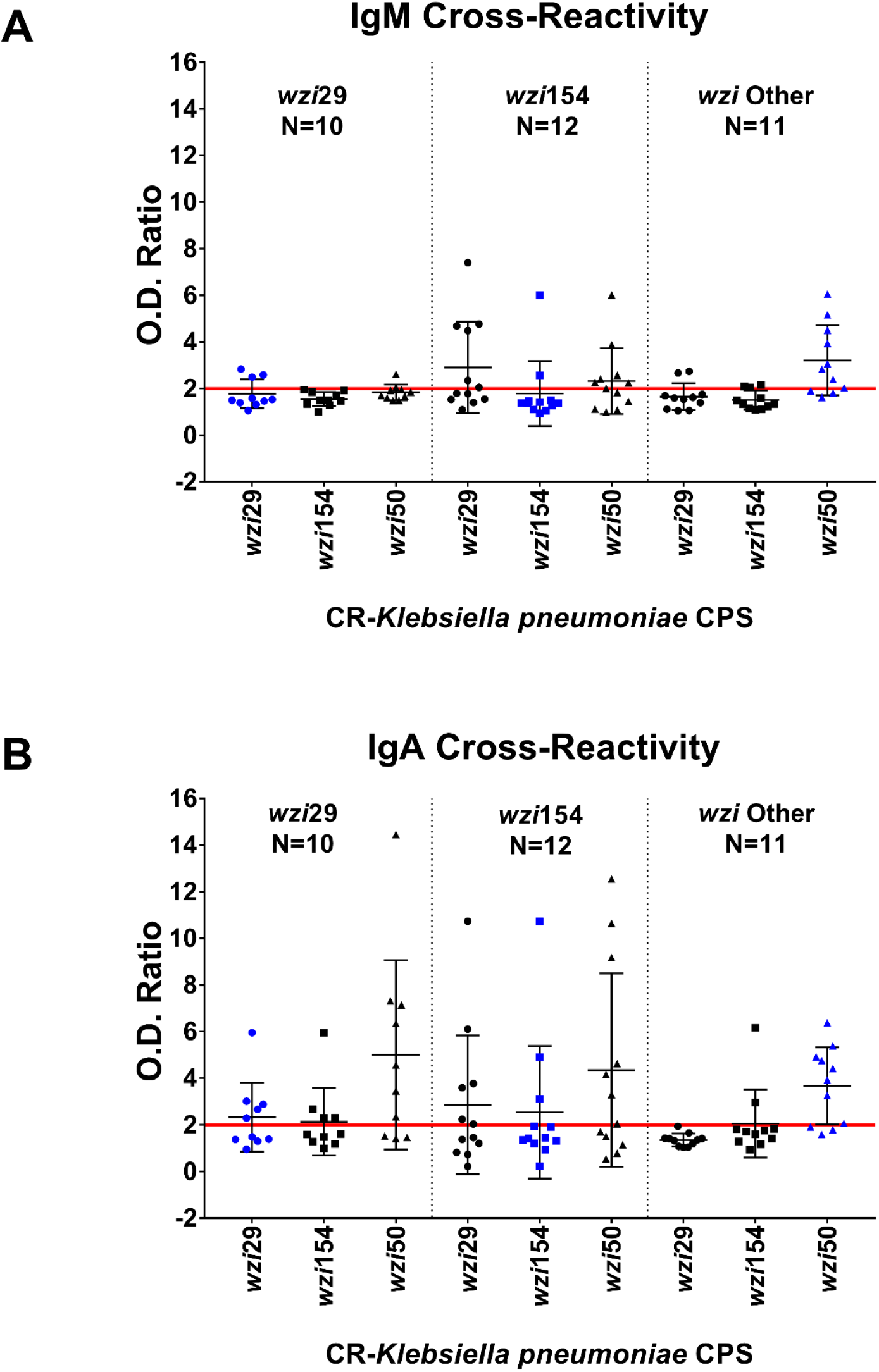
Antibody responses to capsular polysaccharides (CPS) and lipopolysaccharides (LPS) isolated from carbapenem-resistant *K. pneumoniae*. IgM response **(A)** and IgA response **(B)** to *wzi29* (blue dots in the first section); to *wzi154* (blue dots in the middle section); to *wzi50* (blue dots in the last section) was compared in CR-*Kp* patients. O.D. Ratio = (O.D. CPS/O.D. BSA). The positive cut-off is set at Y=2 (red line), the positive serum had the value ≥2 and negative serum is <2; each dot on the scatter plot are individual CR-*Kp* patients’ O.D. ratio and the deviation in O.D. ratios are shown with SD.

**Supplementary Figure 3:**
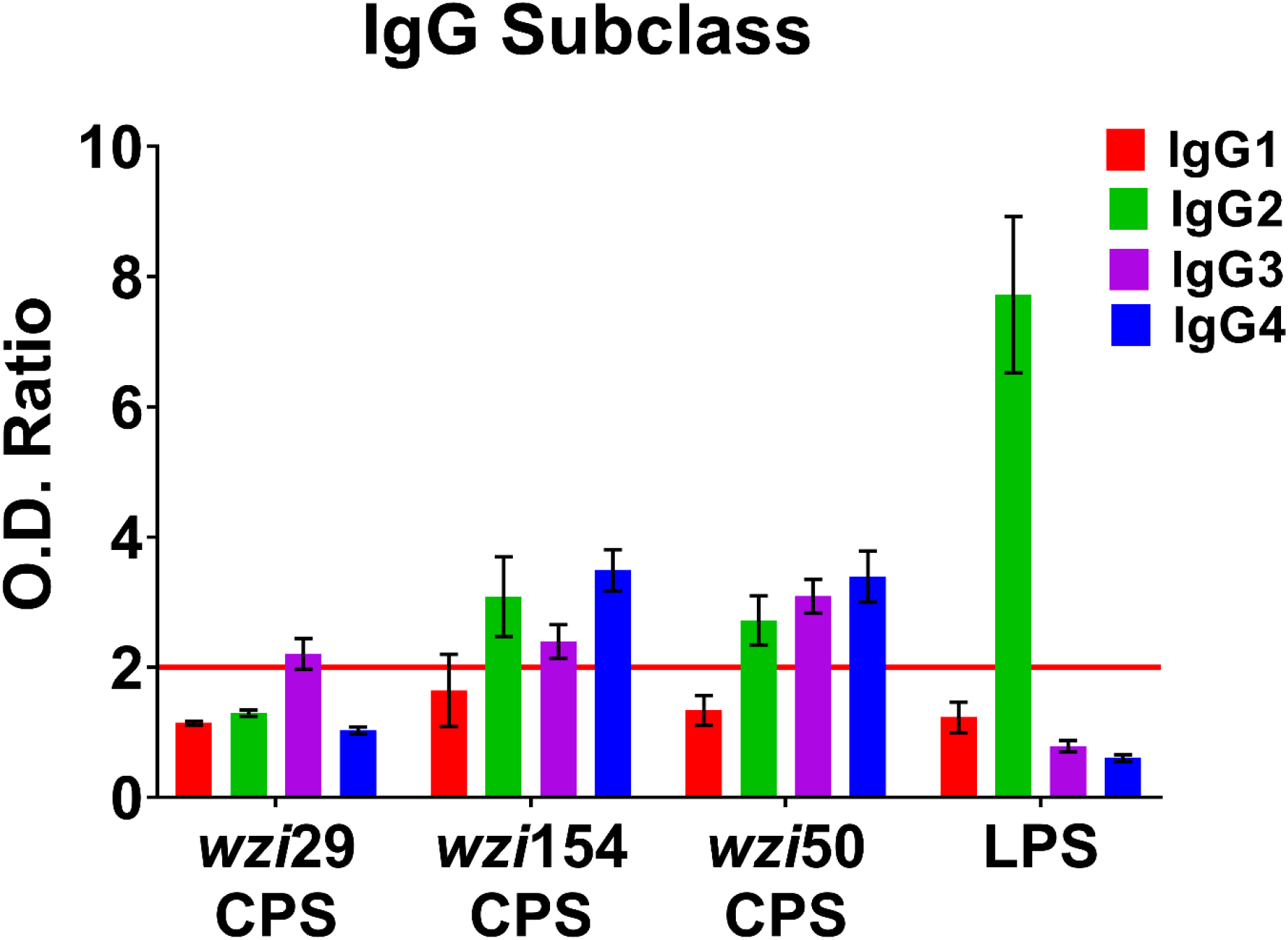
IgG subclass responses to capsular polysaccharides (CPS) and lipopolysaccharides (LPS) in carbapenem-resistant *K. pneumoniae* infected patients (N = 33). IgG1 **(A)**; IgG2 **(B)**; IgG3 **(C)**; IgG4 **(D)**. O.D. Ratio = (O.D. CPS/ O.D. BSA). The positive serum had the value ≥2 and negative serum is <2; bars depict means and SD of three independent experiments, with wells performed in triplicate.

**Supplementary Figure 4:**
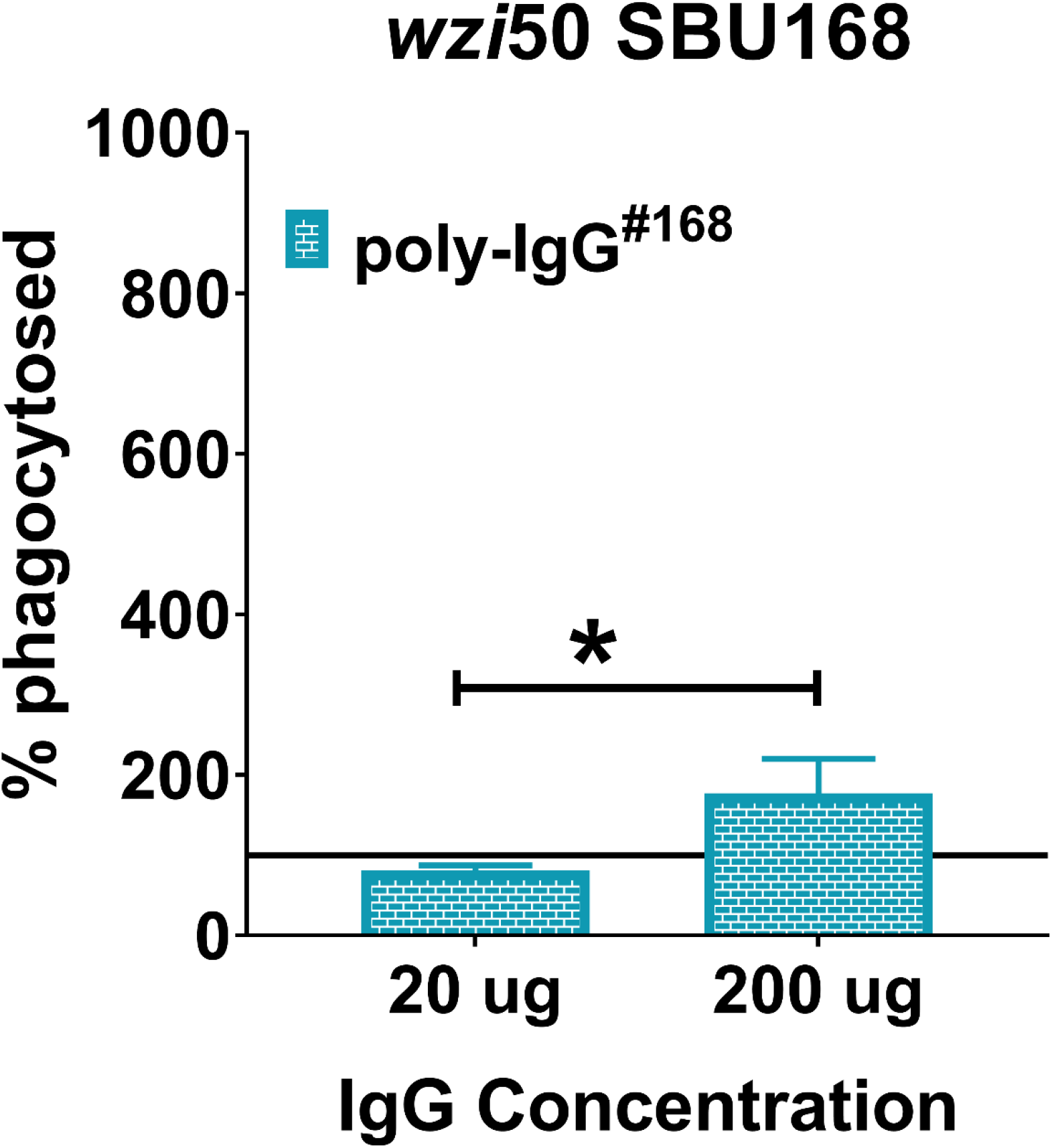
PD-IgGs isolated from patient 168 (*wzi50*) induced opsonophagocytosis of patient-matched SBU168 strain. Phagocytosis cut-off is set at Y=100 (black line) with respect to PBS control, positive phagocytosis had value >100 and no phagocytosis or inhibition of phagocytosis ≤ 100. (** p<0.01, * p<0.05).

## References

1. Kobayashi SD, Porter AR, Dorward DW, Brinkworth AJ, Chen L, Kreiswirth BN, et al. Phagocytosis and Killing of Carbapenem-Resistant ST258 Klebsiella pneumoniae by Human Neutrophils. The Journal of infectious diseases. 2016;213(10):1615–22. doi: 10.1093/infdis/jiw001. PubMed PMID: 26768252; PubMed Central PMCID: PMC4837910.

2. Xu L, Sun X, Ma X. Systematic review and meta-analysis of mortality of patients infected with carbapenem-resistant Klebsiella pneumoniae. Annals of clinical microbiology and antimicrobials. 2017;16(1):18. doi: 10.1186/s12941-017-0191-3. PubMed PMID: 28356109; PubMed Central PMCID: PMC5371217.

3. Organization WH. WHO Publishes Global Priority List of Antibiotic-Resistant Bacteria To Guide Research, Discovery, And Development of New Antibiotics. Geneva: World Health Organization. 2019.

4. van Duin D, Arias CA, Komarow L, Chen L, Hanson BM, Weston G, et al. Molecular and clinical epidemiology of carbapenem-resistant Enterobacterales in the USA (CRACKLE-2): a prospective cohort study. The Lancet Infectious diseases. 2020. doi: 10.1016/S1473-3099(19)30755-8. PubMed PMID: 32151332.

5. Follador R, Heinz E, Wyres KL, Ellington MJ, Kowarik M, Holt KE, et al. The diversity of Klebsiella pneumoniae surface polysaccharides. Microbial genomics. 2016;2(8):e000073. doi: 10.1099/mgen.0.000073. PubMed PMID: 28348868; PubMed Central PMCID: PMC5320592.

6. Brisse S, Passet V, Haugaard AB, Babosan A, Kassis-Chikhani N, Struve C, et al. wzi Gene sequencing, a rapid method for determination of capsular type for Klebsiella strains. Journal of clinical microbiology. 2013;51(12):4073–8. doi: 10.1128/JCM.01924-13. PubMed PMID: 24088853; PubMed Central PMCID: PMC3838100.

7. Deleo FR, Chen L, Porcella SF, Martens CA, Kobayashi SD, Porter AR, et al. Molecular dissection of the evolution of carbapenem-resistant multilocus sequence type 258 Klebsiella pneumoniae. Proceedings of the National Academy of Sciences of the United States of America. 2014;111(13):4988–93. Epub 2014/03/19. doi: 10.1073/pnas.1321364111. PubMed PMID: 24639510; PubMed Central PMCID: PMCPMC3977278.

8. Diago-Navarro E, Chen L, Passet V, Burack S, Ulacia-Hernando A, Kodiyanplakkal RP, et al. Carbapenem-resistant Klebsiella pneumoniae exhibit variability in capsular polysaccharide and capsule associated virulence traits. The Journal of infectious diseases. 2014;210(5):803–13. doi: 10.1093/infdis/jiu157. PubMed PMID: 24634498; PubMed Central PMCID: PMC4432395.

9. Satlin MJ, Chen L, Patel G, Gomez-Simmonds A, Weston G, Kim AC, et al. Multicenter Clinical and Molecular Epidemiological Analysis of Bacteremia Due to Carbapenem-Resistant Enterobacteriaceae (CRE) in the CRE Epicenter of the United States. Antimicrobial agents and chemotherapy. 2017;61(4). doi: 10.1128/AAC.02349-16. PubMed PMID: 28167547; PubMed Central PMCID: PMC5365653.

10. Conte V, Monaco M, Giani T, D’Ancona F, Moro ML, Arena F, et al. Molecular epidemiology of KPC-producing Klebsiella pneumoniae from invasive infections in Italy: increasing diversity with predominance of the ST512 clade II sublineage. J Antimicrob Chemother. 2016;71(12):3386–91. doi: 10.1093/jac/dkw337. PubMed PMID: 27585968.

11. Gomez-Simmonds A, Greenman M, Sullivan SB, Tanner JP, Sowash MG, Whittier S, et al. Population Structure of Klebsiella pneumoniae Causing Bloodstream Infections at a New York City Tertiary Care Hospital: Diversification of Multidrug-Resistant Isolates. J Clin Microbiol. 2015;53(7):2060–7. doi: 10.1128/JCM.03455-14. PubMed PMID: 25878348; PubMed Central PMCID: PMCPMC4473223.

12. van Duin D, Perez F, Rudin SD, Cober E, Hanrahan J, Ziegler J, et al. Surveillance of carbapenem-resistant Klebsiella pneumoniae: tracking molecular epidemiology and outcomes through a regional network. Antimicrob Agents Chemother. 2014;58(7):4035–41. doi: 10.1128/AAC.02636-14. PubMed PMID: 24798270; PubMed Central PMCID: PMCPMC4068524.

13. DeLeo FR, Kobayashi SD, Porter AR, Freedman B, Dorward DW, Chen L, et al. Survival of Carbapenem-Resistant Klebsiella pneumoniae Sequence Type 258 in Human Blood. Antimicrobial agents and chemotherapy. 2017;61(4). doi: 10.1128/AAC.02533-16. PubMed PMID: 28115349; PubMed Central PMCID: PMC5365663.

14. Jansen KU, Anderson AS. The role of vaccines in fighting antimicrobial resistance (AMR). Human vaccines & immunotherapeutics. 2018;14(9):2142–9. doi: 10.1080/21645515.2018.1476814. PubMed PMID: 29787323; PubMed Central PMCID: PMC6183139.

15. Diago-Navarro E, Calatayud-Baselga I, Sun D, Khairallah C, Mann I, Ulacia-Hernando A, et al. Antibody-Based Immunotherapy To Treat and Prevent Infection with Hypervirulent Klebsiella pneumoniae. Clinical and vaccine immunology: CVI. 2017;24(1). doi: 10.1128/CVI.00456-16. PubMed PMID: 27795303; PubMed Central PMCID: PMC5216427.

16. Diago-Navarro E, Motley MP, Ruiz-Perez G, Yu W, Austin J, Seco BMS, et al. Novel, Broadly Reactive Anticapsular Antibodies against Carbapenem-Resistant Klebsiella pneumoniae Protect from Infection. MBio. 2018;9(2). Epub 2018/04/05. doi: 10.1128/mBio.00091-18. PubMed PMID: 29615497; PubMed Central PMCID: PMCPMC5885035.

17. Kobayashi SD, Porter AR, Freedman B, Pandey R, Chen L, Kreiswirth BN, et al. Antibody-Mediated Killing of Carbapenem-Resistant ST258 Klebsiella pneumoniae by Human Neutrophils. mBio. 2018;9(2). doi: 10.1128/mBio.00297-18. PubMed PMID: 29535199; PubMed Central PMCID: PMC5850326.

18. Opoku-Temeng C, Kobayashi SD, DeLeo FR. Klebsiella pneumoniae capsule polysaccharide as a target for therapeutics and vaccines. Computational and structural biotechnology journal. 2019;17:1360–6. doi: 10.1016/j.csbj.2019.09.011. PubMed PMID: 31762959; PubMed Central PMCID: PMC6861629.

19. Malachowa N, Kobayashi SD, Porter AR, Freedman B, Hanley PW, Lovaglio J, et al. Vaccine Protection against Multidrug-Resistant Klebsiella pneumoniae in a Nonhuman Primate Model of Severe Lower Respiratory Tract Infection. mBio. 2019;10(6). doi: 10.1128/mBio.02994-19. PubMed PMID: 31848292; PubMed Central PMCID: PMC6918093.

20. Bloom DE, Black S, Salisbury D, Rappuoli R. Antimicrobial resistance and the role of vaccines. Proceedings of the National Academy of Sciences of the United States of America. 2018;115(51):12868–71. doi: 10.1073/pnas.1717157115. PubMed PMID: 30559204; PubMed Central PMCID: PMC6305009.

21. Rollenske T, Szijarto V, Lukasiewicz J, Guachalla LM, Stojkovic K, Hartl K, et al. Cross-specificity of protective human antibodies against Klebsiella pneumoniae LPS O-antigen. Nature immunology. 2018;19(6):617–24. doi: 10.1038/s41590-018-0106-2. PubMed PMID: 29760533.

22. Calzas C, Lemire P, Auray G, Gerdts V, Gottschalk M, Segura M. Antibody response specific to the capsular polysaccharide is impaired in Streptococcus suis serotype 2-infected animals. Infection and immunity. 2015;83(1):441–53. doi: 10.1128/IAI.02427-14. PubMed PMID: 25385801; PubMed Central PMCID: PMC4288899.

23. Wang Y, Wen Z, Pan X, Briles DE, He Y, Zhang JR. Novel Immunoprotective Proteins of Streptococcus pneumoniae Identified by Opsonophagocytosis Killing Screen. Infection and immunity. 2018;86(9). doi: 10.1128/IAI.00423-18. PubMed PMID: 29891544; PubMed Central PMCID: PMC6105882.

24. Feldman MF, Mayer Bridwell AE, Scott NE, Vinogradov E, McKee SR, Chavez SM, et al. A promising bioconjugate vaccine against hypervirulent Klebsiella pneumoniae. Proceedings of the National Academy of Sciences of the United States of America. 2019;116(37):18655–63. doi: 10.1073/pnas.1907833116. PubMed PMID: 31455739; PubMed Central PMCID: PMC6744904.

25. Campbell WN, Hendrix E, Cryz S, Jr., Cross AS. Immunogenicity of a 24-valent Klebsiella capsular polysaccharide vaccine and an eight-valent Pseudomonas O-polysaccharide conjugate vaccine administered to victims of acute trauma. Clinical infectious diseases: an official publication of the Infectious Diseases Society of America. 1996;23(1):179–81. doi: 10.1093/clinids/23.1.179. PubMed PMID: 8816151.

26. Cryz SJ, Jr., Furer E, Germanier R. Immunization against fatal experimental Klebsiella pneumoniae pneumonia. Infection and immunity. 1986;54(2):403–7. PubMed PMID: 3533779; PubMed Central PMCID: PMC260175.

27. Edelman K. Multiple pyogenic liver abscesses communicating with the biliary tree: treatment by endoscopic stenting and stone removal. The American journal of gastroenterology. 1994;89(11):2070–2. PubMed PMID: 7942740.

28. Kim D, Park BY, Choi MH, Yoon EJ, Lee H, Lee KJ, et al. Antimicrobial resistance and virulence factors of Klebsiella pneumoniae affecting 30 day mortality in patients with bloodstream infection. The Journal of antimicrobial chemotherapy. 2019;74(1):190–9. doi: 10.1093/jac/dky397. PubMed PMID: 30295771.

29. Kubler-Kielb J, Vinogradov E, Ng WI, Maczynska B, Junka A, Bartoszewicz M, et al. The capsular polysaccharide and lipopolysaccharide structures of two carbapenem resistant Klebsiella pneumoniae outbreak isolates. Carbohydrate research. 2013;369:6–9. doi: 10.1016/j.carres.2012.12.018. PubMed PMID: 23360863; PubMed Central PMCID: PMC3594109.

30. Lang AB, Bruderer U, Senyk G, Pitt TL, Larrick JW, Cryz SJ, Jr. Human monoclonal antibodies specific for capsular polysaccharides of Klebsiella recognize clusters of multiple serotypes. Journal of immunology. 1991;146(9):3160–4. PubMed PMID: 2016541.

31. Smith K, Muther JJ, Duke AL, McKee E, Zheng NY, Wilson PC, et al. Fully human monoclonal antibodies from antibody secreting cells after vaccination with Pneumovax(R)23 are serotype specific and facilitate opsonophagocytosis. Immunobiology. 2013;218(5):745–54. doi: 10.1016/j.imbio.2012.08.278. PubMed PMID: 23084371; PubMed Central PMCID: PMC3556204.

32. Seeberger PH, Pereira CL, Khan N, Xiao G, Diago-Navarro E, Reppe K, et al. A Semi-Synthetic Glycoconjugate Vaccine Candidate for Carbapenem-Resistant Klebsiella pneumoniae. Angewandte Chemie. 2017;56(45):13973–8. doi: 10.1002/anie.201700964. PubMed PMID: 28815890; PubMed Central PMCID: PMC5819008.

33. Yokota S, Amano K, Hayashi S, Kubota T, Fujii N, Yokochi T. Human antibody response to Helicobacter pylori lipopolysaccharide: presence of an immunodominant epitope in the polysaccharide chain of lipopolysaccharide. Infection and immunity. 1998;66(6):3006–11. PubMed PMID: 9596783; PubMed Central PMCID: PMC108305.

34. Chen T, Blanc C, Liu Y, Ishida E, Singer S, Xu J, et al. Capsular glycan recognition provides antibody-mediated immunity against tuberculosis. The Journal of clinical investigation. 2020;130(4):1808–22. doi: 10.1172/JCI128459. PubMed PMID: 31935198; PubMed Central PMCID: PMC7108924.

35. Clements A, Jenney AW, Farn JL, Brown LE, Deliyannis G, Hartland EL, et al. Targeting subcapsular antigens for prevention of Klebsiella pneumoniae infections. Vaccine. 2008;26(44):5649–53. doi: 10.1016/j.vaccine.2008.07.100. PubMed PMID: 18725260.

36. Hegerle N, Choi M, Sinclair J, Amin MN, Ollivault-Shiflett M, Curtis B, et al. Development of a broad spectrum glycoconjugate vaccine to prevent wound and disseminated infections with Klebsiella pneumoniae and Pseudomonas aeruginosa. PloS one. 2018;13(9):e0203143. doi: 10.1371/journal.pone.0203143. PubMed PMID: 30188914; PubMed Central PMCID: PMC6126813 and patent applications (U.S. Patent Application No. 15/842,068, European Patent Application No. 15 841 265.0) describing Klebsiella reagent strains and conjugates of Klebsiella OPS with Pseudomonas flagellin proteins. AL and MOS are employees of Fina Biosolutions. AL is also a patent holder for the aminooxy method used for conjugation (WO2005072778 A2). This does not alter our adherence to PLOS ONE policies on sharing data and materials.

37. Tzouvelekis LS, Miriagou V, Kotsakis SD, Spyridopoulou K, Athanasiou E, Karagouni E, et al. KPC-producing, multidrug-resistant Klebsiella pneumoniae sequence type 258 as a typical opportunistic pathogen. Antimicrobial agents and chemotherapy. 2013;57(10):5144–6. doi: 10.1128/AAC.01052-13. PubMed PMID: 23856769; PubMed Central PMCID: PMC3811402.

38. van Duin D, Paterson DL. Multidrug-Resistant Bacteria in the Community: Trends and Lessons Learned. Infectious disease clinics of North America. 2016;30(2):377–90. doi: 10.1016/j.idc.2016.02.004. PubMed PMID: 27208764; PubMed Central PMCID: PMC5314345.

39. Martin RM, Bachman MA. Colonization, Infection, and the Accessory Genome of Klebsiella pneumoniae. Frontiers in cellular and infection microbiology. 2018;8:4. doi: 10.3389/fcimb.2018.00004. PubMed PMID: 29404282; PubMed Central PMCID: PMC5786545.

40. Hicks LA, Harrison LH, Flannery B, Hadler JL, Schaffner W, Craig AS, et al. Incidence of pneumococcal disease due to non-pneumococcal conjugate vaccine (PCV7) serotypes in the United States during the era of widespread PCV7 vaccination, 1998-2004. The Journal of infectious diseases. 2007;196(9):1346–54. doi: 10.1086/521626. PubMed PMID: 17922399.

41. Singleton RJ, Hennessy TW, Bulkow LR, Hammitt LL, Zulz T, Hurlburt DA, et al. Invasive pneumococcal disease caused by nonvaccine serotypes among alaska native children with high levels of 7-valent pneumococcal conjugate vaccine coverage. Jama. 2007;297(16):1784–92. doi: 10.1001/jama.297.16.1784. PubMed PMID: 17456820.

42. Brito LA, Singh M. Acceptable levels of endotoxin in vaccine formulations during preclinical research. Journal of pharmaceutical sciences. 2011;100(1):34–7. doi: 10.1002/jps.22267. PubMed PMID: 20575063.

43. Wu MF, Yang CY, Lin TL, Wang JT, Yang FL, Wu SH, et al. Humoral immunity against capsule polysaccharide protects the host from magA+ Klebsiella pneumoniae-induced lethal disease by evading Toll-like receptor 4 signaling. Infection and immunity. 2009;77(2):615–21. doi: 10.1128/IAI.00931-08. PubMed PMID: 19015249; PubMed Central PMCID: PMC2632026.

44. Diaz Romero J, Outschoorn I. Selective biotinylation of Neisseria meningitidis group B capsular polysaccharide and application in an improved ELISA for the detection of specific antibodies. Journal of immunological methods. 1993;160(1):35–47. doi: 10.1016/0022-1759(93)90006-s. PubMed PMID: 8450238.

